# Golden magic: RSH enzymes for (p)ppGpp metabolism in the diatom *Phaeodactylum tricornutum*

**DOI:** 10.1101/487603

**Authors:** L Avilan, C Puppo, A Villain, E Bouveret, B Menand, B Field, B Gontero

**Affiliations:** Aix Marseille Univ CNRS, BIP, UMR 7281, IMM FR 3479, 31 Chemin Joseph Aiguier, Marseille, France F-13009; Aix Marseille Univ CNRS, IGS; UMR 7256, IMM FR 3479, Marseille, France F-13009; Stress Adaptation and Metabolism in Enterobacteriae group, Department of Microbiology, Institut Pasteur, Paris 75015, France; Aix Marseille Univ, CEA, CNRS, BIAM, Laboratoire de génétique et biophysique des plantes, Marseille, France F-13009

**Keywords:** Chloroplast, diatom, *Phaeodactylum tricornutum*, plastid, (p)ppGpp, RSH

## Abstract

The nucleotides guanosine tetraphosphate and pentaphosphate (together known as (p)ppGpp or magic spot) are produced in plant plastids from GDP/GTP and ATP by RelA-SpoT homologue (RSH) enzymes. In the model plant Arabidopsis (p)ppGpp regulates chloroplast transcription and translation to affect growth, and is also implicated in acclimation to stress. However, little is known about (p)ppGpp metabolism or its evolution in other photosynthetic eukaryotes. Here we studied (p)ppGpp metabolism in the golden-coloured marine diatom *Phaeodactylum tricornutum*. We identified three expressed *RSH* genes in the *P. tricornutum* genome, and determined the enzymatic activity of the corresponding enzymes by heterologous expression in bacteria. We showed that two *P. tricornutum* RSH are (p)ppGpp synthetases, despite substitution of a residue within the active site believed critical for activity, and that the third RSH is a bifunctional (p)ppGpp synthetase and hydrolase, the first of its kind demonstrated in a photosynthetic eukaryote. A broad phylogenetic analysis then showed that diatom RSH belong to novel algal RSH clades. Together our work significantly expands the horizons of (p)ppGpp signalling in the photosynthetic eukaryotes by demonstrating an unexpected functional, structural and evolutionary diversity in RSH enzymes from organisms with plastids derived from red algae.

**Highlight:** We discover RSH enzymes for (p)ppGpp metabolism in the diatom *Phaeodactylum tricornutum* and show that they have surprising functional and structural features, and belong to novel red-plastid lineage RSH clades.

## Introduction

Plastids, the defining feature of photosynthetic eukaryotes, arose from the endosymbiosis of a cyanobacterium more than one billion years ago (Ponce-Toledo *et al.*, 2017). Massive cyanobacterial gene loss occurred following endosymbiosis, along with the transfer of genes to the nuclear genome of the eukaryotic host. Plastids now retain a small genome that encodes housekeeping and photosynthetic proteins, as well as a bacteria-like gene expression machinery. Bacteria-like regulatory systems that may be involved in acclimation to environmental perturbation are also present in plastids (Puthiyaveetil *et al.*, 2008; Field, 2018). One of these regulatory-systems is mediated by the nucleotides guanosine tetraphosphate and pentaphosphate (referred to as (p)ppGpp hereafter) whose levels are controlled by the antagonistic action of RelA-SpoT homologues (RSH). In bacteria, (p)ppGpp was originally identified as a magic spot on thin-layer chromatography plates in the 1960s (Cashel & Gallant, 1969). Now bacterial (p)ppGpp signalling is well characterised: (p)ppGpp accumulates in response to stress to reduce proliferation and activates acclimatory pathways by targeting enzymes involved in transcription, translation, and replication (Dalebroux & Swanson, 2012; Hauryliuk *et al.*, 2015).

In the photosynthetic eukaryotes, (p)ppGpp was discovered more recently, and (p)ppGpp signalling has principally been studied in flowering plants. Although (p)ppGpp accumulates in response to various different stresses (Takahashi *et al.*, 2004; Ihara *et al.*, 2015), the actual role of (p)ppGpp during stress acclimation is not yet clear. However, (p)ppGpp is known to act as a potent inhibitor of plastid gene expression *in vivo* (Maekawa *et al.*, 2015; Yamburenko *et al.*, 2015; Sugliani *et al.*, 2016), and altering the capacity of a plant to make (p)ppGpp influences photosynthetic capacity, growth and development (Maekawa *et al.*, 2015; Sugliani *et al.*, 2016). Notably, (p)ppGpp appears to be important for regulating the equilibrium between the plastidic and nucleocytoplasmic compartments of the plant cell. Phylogenetic studies support the existence of at least three conserved families of plastid-targeted RSH enzymes in land plants named RSH1, RSH2/3 and RSH4 (Atkinson *et al.*, 2011). The model flowering plant *Arabidopsis thaliana*, where plant (p)ppGpp homeostasis is the most well understood, possesses representatives from each of these families: RSH1 that lacks (p)ppGpp synthetase activity and appears to function as the major (p)ppGpp hydrolase (Sugliani *et al.*, 2016), the closely related RSH2 and RSH3 that appear to act as the major (p)ppGpp synthetases (Mizusawa *et al.*, 2008; Maekawa *et al.*, 2015; Sugliani *et al.*, 2016), and a calcium- activated RSH (CRSH) from the RSH4 family that possesses a C-terminal EF-hand domain implicated in calcium binding, and has calcium-dependent (p)ppGpp synthesis activity *in vitro* (Masuda *et al.*, 2008).

Plastids are the result of primary endosymbiosis, where a photosynthetic bacterium was engulfed by a eukaryote. Today several primary plastid lineages can be identified which are thought to share common ancestry. Plants and green algae (together known as the Viridiplantae) possess primary plastids of the green lineage. The Viridiplantae have two sister groups with primary plastids, the red algae (Rhodophyta) and glaucophytes (Glaucophyta). Primary plastids from the green and red lineages have also been transferred and mixed in new eukaryotic hosts to result in the astonishing range of plastid diversity that can be observed in nature (Archibald, 2015). Plastid-targeted RSH enzymes from the RSH1, RSH2/3 and RSH4 families have been identified in many green and red algae, yet the metabolism and functions of (p)ppGpp in these organisms have barely been investigated (Atkinson *et al.*, 2011; Ito *et al.*, 2017; Field, 2018). One exception is a recent report on the red alga *Cyanidioschyzon merolae*, where CmRSH4b, a member of the RSH4 family, was shown to possess (p)ppGpp synthetase activity (Imamura *et al.*, 2018). Interestingly, the inducible expression of CmRSH4b in *C. merolae* results in a reduction in plastid size and rRNA transcription in a similar manner to the expression of a (p)ppGpp synthetase in Arabidopsis (Sugliani *et al.*, 2016).

Diatoms (Bacillariophyceae) are a group of golden-coloured microalgae that contain complex plastids that originate from the secondary endosymbiosis of a red alga, with the addition of nuclear-encoded plastid-targeted green algal proteins left over from a previous endosymbiosis (Dorrell & Smith, 2011; Dorrell *et al.*, 2017). Diatoms are the predominant photosynthetic eukaryote in the oceans, where they account for around 40% of net primary production (Malviya *et al.*, 2016). Therefore, understanding (p)ppGpp synthesis in diatoms, where it is likely to regulate photosynthetic capacity (Maekawa *et al.*, 2015; Sugliani *et al.*, 2016), and play roles in diatom lifestyle, is of particular importance. To tackle this issue, we investigate (p)ppGpp metabolism in the model pennate marine diatom *Phaeodactylum tricornutum*. We first identified expressed *RSH* genes in the *P. tricornutum* genome and then determined if their gene products have (p)ppGpp synthetase / hydrolase activity by complementation of *Escherichia coli* (p)ppGpp biosynthesis mutants. Then, we place the structural and catalytic features of *P. tricornutum* RSH enzymes into an evolutionary context. Altogether our study advances our previously poor understanding of (p)ppGpp metabolism in the complex plastids of diatoms.

## Materials and Methods

### Cloning of *P. tricornutum* RSH

Genomic DNA from *P. tricornutum* Bohlin (strain number CCAP 1052/1A from Culture Collection of Algae and Protozoa, CCAP, Scottish Marine Institute) was isolated from cells grown in late-exponential phase, using an optimized method for diatoms (Puppo *et al.*, 2017; Kojadinovic-Sirinelli *et al.*, 2018). Sequences for the *P. tricornutum* RSH proteins: PtRSH1, PtRSH4a and PtRSH4b (accession codes: 11099, 7629 and 3397, respectively) were retrieved from Joint Genome Initiative (JGI) database (https://genome.jgi.doe.gov/Phatr2/Phatr2.home.html). The amino acid sequences of PtRSH1 and PtRSH4a reported in JGI were partial, so a manual search based on nucleotide sequence was performed to obtain the full-length amino acid sequences. The three PtRSH sequences are given in Fig. S1. The genes coding for the mature proteins without the bipartite chloroplast targeting peptide were amplified by PCR using the primers listed in Table S1. The theoretical transit peptide cleavage site was estimated from (Huesgen *et al.*, 2013) and by using alignments with plant RSH enzymes. Primers were designed to have 20-bp flanking regions homologous to the plasmid (pBAD24) each side of the *Nco*I and *Hind*III restriction sites. The plasmid construction was carried out using the Sequence and Ligation Independent Cloning (SLIC) method (Jeong *et al.*, 2012). PCR was performed using genomic DNA of *P tricornutum*, the specified primers and Q5 DNA polymerase (New England Biolabs). Since the gene encoding PtRSH1 has an intron, the complete cloned gene was used as a template for the amplification of the two exons using the primers specified in Table S1. The final construct was assembled in the pBAD24 plasmid using SLIC cloning (Jeong *et al.*, 2012). All constructs were verified by sequencing (GATC Biotech).

### RSH activity assays

To test (p)ppGpp synthetase activity, pBAD24 plasmids containing the *PtRSH* genes, SpoT or without a coding-sequence (empty) were transformed into *E. coli* strain EB425 (MG1655 Δ*relA* Δ*spoT*) (Wahl *et al.*, 2011) and grown at 37°C on M9 minimal medium agar plates without amino acids and in the presence or absence of 0.1% arabinose. To test (p)ppGpp hydrolase activity, the same plasmids were transformed into *E. coli* strain EB544 (MG1655 Δ*relA spoT*203) (My *et al.*, 2013). Pre-cultures from independent colonies for each replicate were diluted in 50 ml Luria Bertani medium containing ampicillin (100 μg mL^−1^) and arabinose (0.1% w/v). Growth was performed at 37°C under agitation at 175 rpm and optical density was measured at 600 nm every 30 min.

### Prediction of plant and algal RSH enzyme structures

Iterative Threading ASSEmbly Refinement (iTASSER)(Yang *et al.*, 2015) was used to predict the structure and ligands of the synthetase and hydrolase domain containing region of Arabidopsis and *P. tricornutum* RSH enzymes (default settings). The synthetase hydrolase regions were chosen by selection of the regions corresponding most closely to the synthetase region of *Bacillus subtilis* SAS1: AtRSH1, residues 347-542; AtRSH3, residues 404-559; PtRSH1, residues 403-584; PtRSH4a, residues 352-537; PtRSH4b, residues 432-631. The *B. subtilis* SAS1 crystal structure (9decA) was then manually selected in iTASSER for modelling. Results were visualized in PyMOL (Version 2.0 Schrödinger, LLC).

### Phylogenetic analysis of RSH proteins from diatoms and other photosynthetic organisms

Using *P. tricornutum* RSH proteins as queries, a set of homologous proteins was built using public data at the National Center for Biotechnology Information (NCBI) and JGI, as well as individual genome projects : *Synedra acus* (http://www.lin.irk.ru/sacus/index.php?r=site/page&view=downloads&lang=en), *Cyclotella cryptica* (http://genomes.mcdb.ucla.edu/Cyclotella/download.html), *Pseudo nitzschia multistriata* (https://zenodo.org/record/495408#.W1sOJsIyXIU) and *Asterionella formosa* (unpublished data, A. Villain, personal communication). RSH from similar plant and green algal species to those selected by Ito et al., (2017) were used. RSH proteins were selected only from fully sequenced genomes, and all predicted RSH per genome were used in the subsequent analysis. Two datasets were built: a ‘whole dataset’ containing all RSH sequences, and a ‘RSH1 specific set’ including only RSH1 sequences. Multiple-sequence alignments of homologous proteins were performed separately on both datasets using MAFFT v7.402 (Katoh *et al.*, 2017) with option –auto. Initial alignment of the ‘whole dataset’ was trimmed to include only the hydrolase and synthetase domains, by selecting columns 26 to 343 from *E. coli* SpoT, while full-length sequences were kept in the ‘RSH1 specific’ alignment. Alignment columns containing gaps in >30% sequences were then removed from both alignments (Supplementary dataset), and maximum likelihood phylogenetic reconstructions were performed with FastTree version 2.1.10 (Price *et al.*, 2010) with options -slow -mlacc 3 -wag -gamma. Protein domain architecture was analyzed by CD-search algorithm of NCBI with default parameters and an E-value threshold of 0.01 (Marchler-Bauer *et al.*, 2015).

## Results

### The nuclear genome of the model diatom *Phaeodactfyum tricornutum* encodes three plastid-targeted RSH enzymes

We inspected the *P. tricornutum* genome for the presence of *RSH* genes using BLAST and identified three: *PtRSH1*, *PtRSH4a* and *PtRSH4b*. *PtRSH1* is the only *RSH* gene that contains an intron. We analysed the predicted protein sequences derived from the three genes and identified a number of conserved domains that are typical of RSH enzymes (Fig. 1A). The N-terminus of all the three RSH also possesses a domain with strong resemblance to a bipartite signal peptide, which is necessary for the transport of nuclear-encoded proteins to the plastid in diatoms according to Gruber et al. (2015). Analysis of amino acid frequency around the potential cleavage site in each sequence supported this assignment, and suggesting that all three RSH are plastid targeted (Fig 1B).

**Fig. 1.**
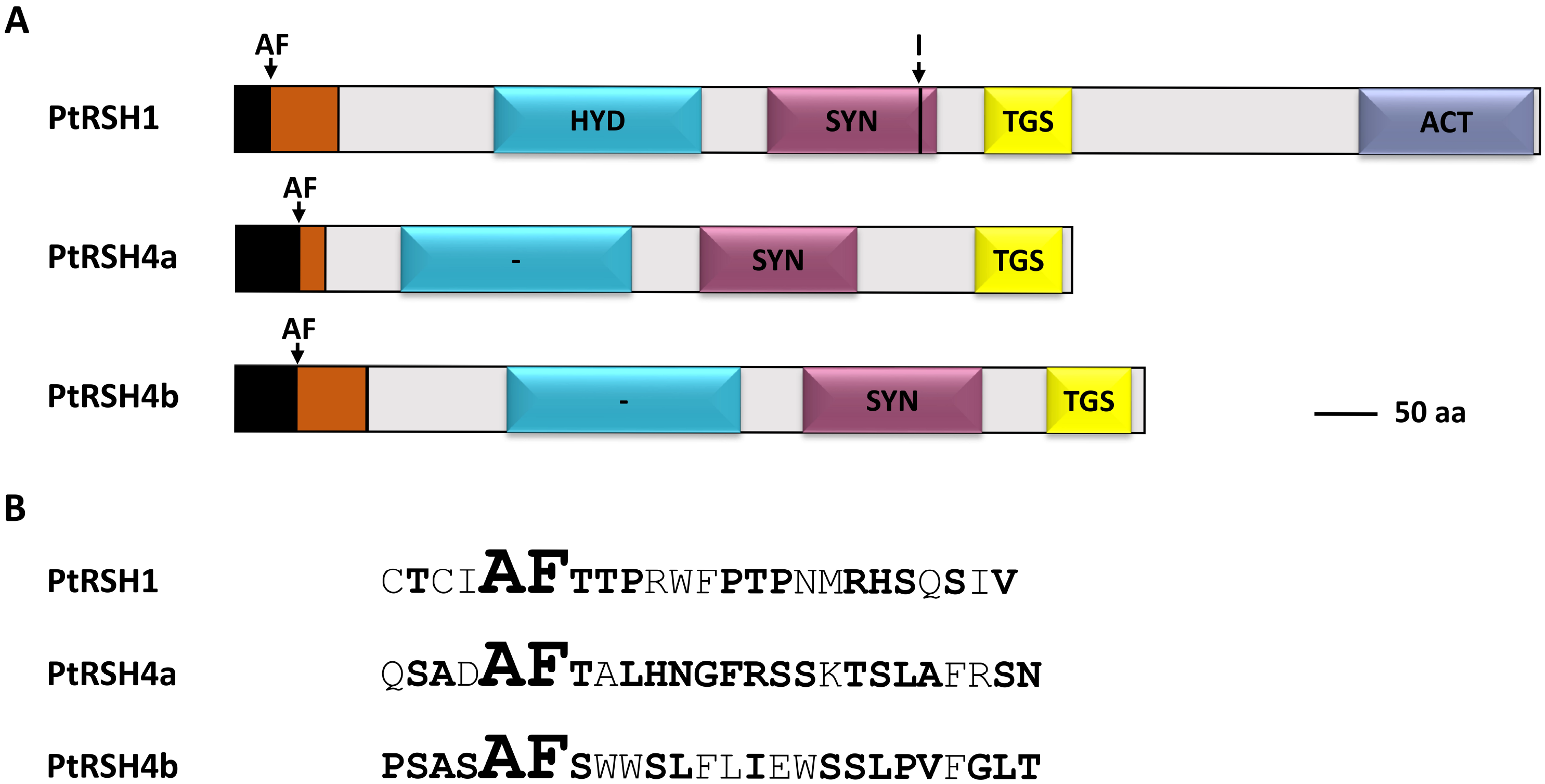
Primary structure of RSH from *P. tricornutum*. (A) Schematic representation of domains found in *P. tricornutum* PtRSH1, PtRSH4a and PtRSH4b (JGI gene accession numbers 11099, 7629 and 33947): (p)ppGpp hydrolase (HYD), (p)ppGpp synthetase (SYN), Threonyl-tRNA synthetase GTPase Spot (TGS) and Aspartate kinase, Chorismate mutase, TyrA (ACT). The bipartite peptide for targeting to the chloroplast is shown. Arrows indicate the cleavage site (AF) of the putative chloroplast signal peptide and the intron position (I) on the corresponding gene. (B) Sequence surrounding the cleavage site (AF) of the putative chloroplast signal peptides of *P. tricornutum* RSH. Bold residues indicate greater than 0.25 frequency of amino acid occurrence (Gruber *et al.*, 2015).

PtRSH4a and PtRSH4b bear (p)ppGpp hydrolase and synthetase domains that show signs of catalytic inactivation (Fig S2). The (p)ppGpp hydrolase domains are divergent compared to the hydrolase domains of known (p)ppGpp hydrolases, and lack many residues critical for hydrolase activity (Steinchen & Bange, 2016). While PtRSH4a and PtRSH4b contain domains with strong homology to bacterial (p)ppGpp synthetase domains, a glycine residue corresponding to G240 in *Streptococcus equisimilis* Rel (Relseq) and previously shown to be essential for synthetase activity in bacterial RSH (Wendrich & Marahiel, 1997; Hogg *et al.*, 2004) is substituted by an alanine (PtRSH4a) or a serine residue (PtRSH4b). Substitution of this glycine residue in the RSH enzymes of land plants has also been associated with the loss of (p)ppGpp synthetase activity (Masuda *et al.*, 2008; Sugliani *et al.*, 2016). In contrast, PtRSH1 bears well conserved (p)ppGpp hydrolase and synthetase domains. This analysis suggests that PtRSH1 may be a bifunctional enzyme, unlike the monofunctional RSH that have so far been identified in plants and green and red algae.

In addition to the N-terminal catalytic domains, all the three PtRSH also possess C-terminal regulatory domains, as observed in bacterial RSH enzymes. PtRSH1, PtRSH4a and PtRSH4b have a TGS domain, named after the proteins where it occurs: Threonyl-tRNA synthetase, GTPase, SpoT and RelA proteins. PtRSH1 has, in addition, an Aspartate kinase, Chorismate mutase, TyrA (ACT) domain, a conserved regulatory fold found in enzymes regulated by small metabolites. The TGS and ACT domains have both been implicated in protein-protein interactions in bacterial RSH enzymes (Wout *et al.*, 2004; Battesti & Bouveret, 2006; Ronneau *et al.*, 2018).

We then analyzed *P. tricornutum* expressed sequence tag (EST) databases (Scala *et al.*, 2002; Bowler *et al.*, 2008) to determine whether the genes encoding PtRSH are expressed. *PtRSH1, PtRSH4a* and *PtRSH4b* are indeed expressed in standard growth conditions (Table S2). *PtRSH4a* had the greatest number of ESTs indicating higher expression levels, and *PtRSH4a* EST abundance increased in response to a number of stresses including high levels of 2E, 4E decadienal, a volatile oxylipin associated with diatom stress signalling (Vardi *et al.*, 2006)(Table S2). This may suggest that diatom *RSH* genes are involved in defence responses, as previously observed for plant *RSH* (van der Biezen *et al.*, 2000; Givens *et al.*, 2004; Kim *et al.*, 2009; Abdelkefi *et al.*, 2018). Furthermore, recent transcriptome experiments show that *PtRSH1* and *PtRSH4b* are induced rapidly after the onset of nitrogen starvation and in response to incubation in darkness for eight hours (Matthijs *et al.*, 2016; Matthijs *et al.*, 2017).

### Catalytic activities of the *P. tricornutum* RSH enzymes

Next, we investigated the potential catalytic activities of PtRSH1, PtRSH4a and PtRSH4b by expressing the predicted mature proteins in *E. coli* strains where the endogenous RSH genes, *relA* and *spoT*, are mutated.

To assay (p)ppGpp synthetase activity, we expressed the *P. tricornutum* RSH enzymes in a Δ*relA* Δ*spoT* mutant. This mutant is unable to grow on minimal medium, and growth can be restored by complementation using a plasmid expressing a functional (p)ppGpp synthetase. The expression of PtRSH1, PtRSH4a and PtRSH4b restored growth to the Δ*relA* Δ*spoT* mutant on minimal medium, as did expression of the positive control *spoT* gene from *E. coli* (Fig. 2A). Thus, all three PtRSH enzymes have (p)ppGpp synthetase activity.

**Fig. 2.**
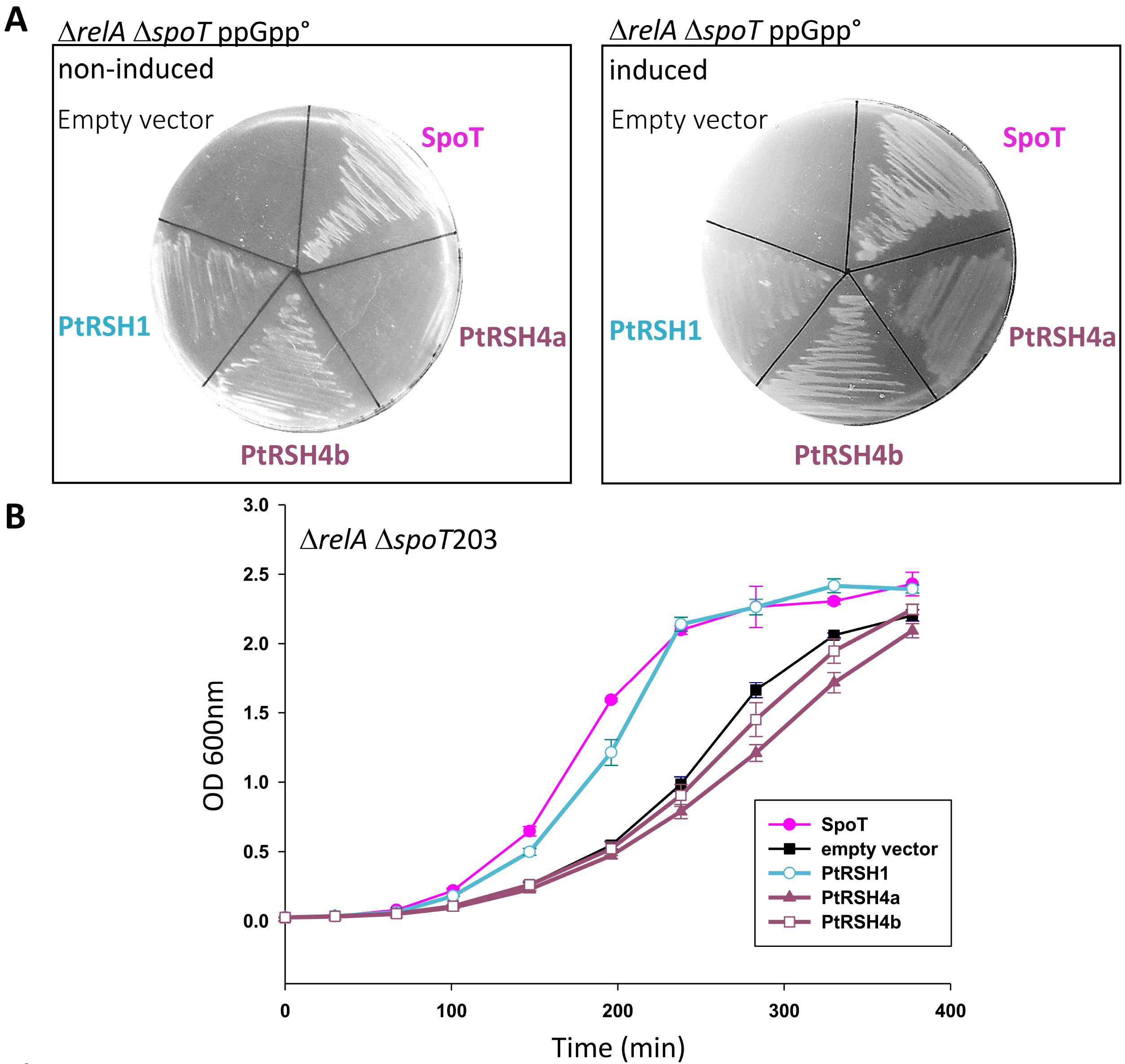
Catalytic activity of *P. tricornutum* RSH. (A) The coding sequences for mature PtRSH1, PtRH4a and PtRSH4b were introduced into the pBAD24 plasmid in MG1655 ΔrelA ΔspoT. Cells were incubated at 37°C on M9 minimal agar medium without amino acids in the absence or presence of the inducer arabinose. Empty vector and pBAD24-Spot from *E.coli* were included as controls. Complementation was also observed without induction, presumably due to leaky expression from pBAD24. (B) Growth curves of MG1655 ΔrelA spoT203 containing coding sequences for the indicated enzymes in the plasmid pBAD24. The values are means ± SD of three biological replicates.

We then used an assay that we recently developed to detect (p)ppGpp hydrolase activity using the Δ*relA spoT*203 mutant strain (Sugliani *et al.*, 2016). In this mutant strain SpoT is defective for (p)ppGpp hydrolase activity while retaining (p)ppGpp synthetase activity, which results in the constitutive over-accumulation of (p)ppGpp and a slow growth phenotype. Normal growth can be restored by expression of an enzyme with (p)ppGpp hydrolase activity such as SpoT. We found that the expression of the diatom PtRSH1 rescued the growth of this mutant in a similar fashion to expression of the positive control *spoT* (Fig. 2B). In contrast, expression of PtRSH4a and PtRSH4b in the Δ*relA spoT*203 mutant strain did not restore normal growth, and we even observed slower growth, perhaps due to enhanced (p)ppGpp biosynthesis. The absence of detectable (p)ppGpp hydrolase activity in PtRSH4a and PtRSH4b is consistent with the divergent (p)ppGpp hydrolase domains in these enzymes that lack many of the residues essential for activity. All together, these results demonstrate that PtRSH4a and PtRSH4b are exclusive (p)ppGpp synthetase enzymes, and that PtRSH1 is a bifunctional RSH enzyme with (p)ppGpp synthetase and hydrolase activity.

### Active site modelling of *P. tricornutum* RSH enzymes

All three RSH from *P. tricornutum* have unusual characteristics. As shown above, PtRSH1 is a bifunctional (p)ppGpp synthetase and hydrolase, a class of enzyme common in bacteria but not yet described in plants or algae. Furthermore, PtRSH4a and PtRSH4b are (p)ppGpp synthetases, despite the lack of a conserved glycine residue previously shown to be critical for (p)ppGpp synthesis as discussed above (Wendrich & Marahiel, 1997; Hogg *et al.*, 2004)(Fig. 3A). Therefore, to gain functional insights into these characteristics, we modelled the structures of the synthetase domains of the three *P. tricornutum* RSH enzymes using I-TASSER (Yang *et al.*, 2015)(Fig. 3B). For the three PtRSH enzymes as well as for CRSH and RSH1 from *A. thaliana* we obtained models that matched the crystal structure of Small Alarmone Synthase 1 (SAS1) from *Bacillus subtilis* bound to the non-hydrolysable ATP analog α,β-methyleneadenosine 5′-triphosphate (AMPCPP) (Steinchen *et al.*, 2015). Within the active site of SAS1 the conserved glycine residue G45 (corresponding to Relseq G240) is positioned on a beta strand neighbouring R46, a residue involved in binding AMPCPP in cooperation with a number of other residues (Fig. 3A, 3B). In PtRSH4a and PtRSH4b the side-chains of the non-conserved residues corresponding to G45 did not greatly affect the active site structure, and their side chains projected away from the active site (Fig. 3B). In both cases, the neighbouring arginine residue corresponding to SAS1 R46 was still predicted to bind AMPCPP. Interestingly, in the residues corresponding to R46 the arginine side-chain took several different orientations in AtRSH3, PtRSH4a, PtRSH4b and PtRSH1, all of which appear to aid in retaining the adenine and ribose moieties of AMPCPP.

**Fig. 3.**
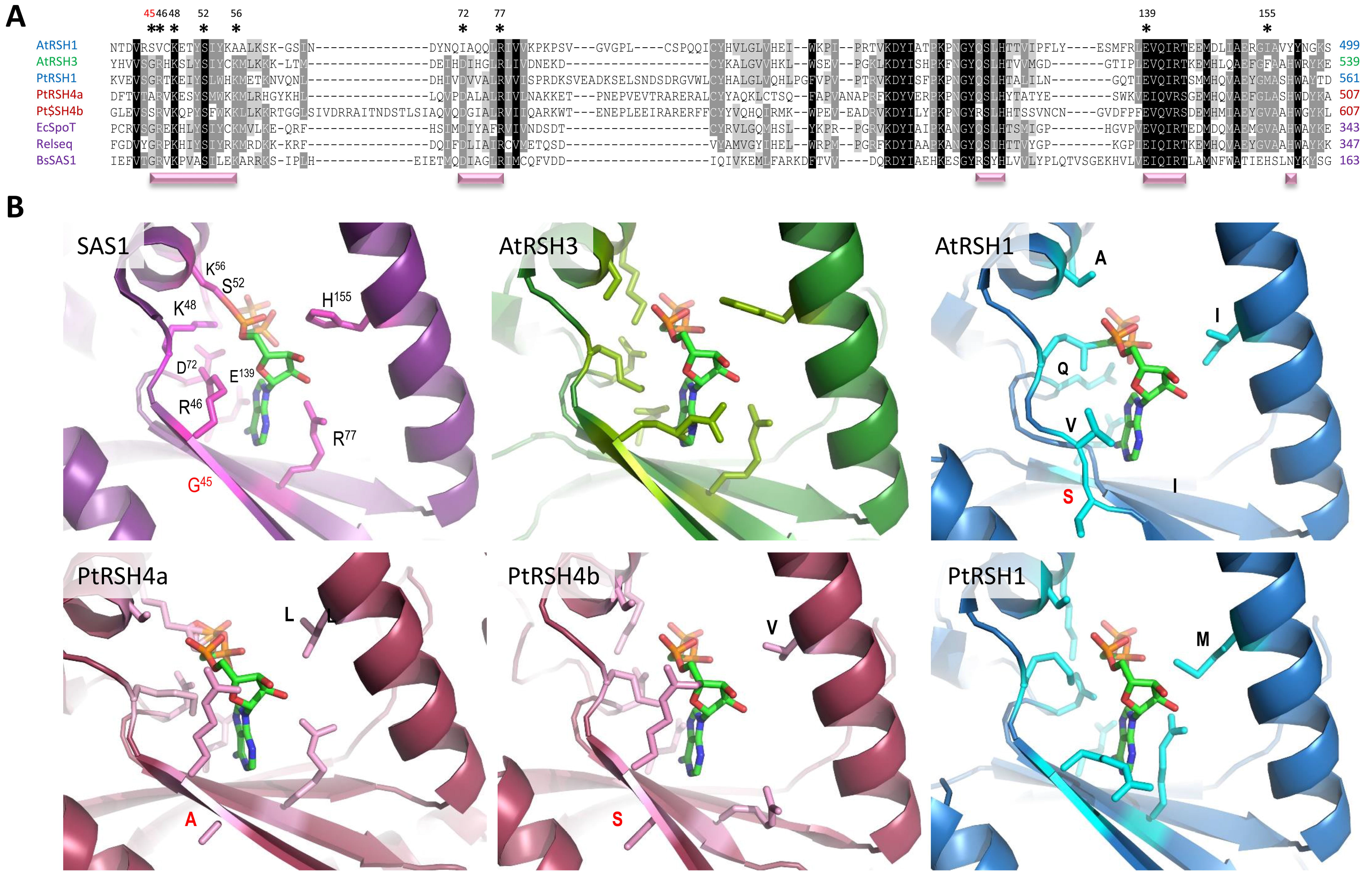
Comparative modelling of the synthetase domain of *P. tricornutum* RSH. (A) Amino acid alignments for the (p)ppGpp synthetase domain of the *B. subtilis* SAS1 and *P. tricornutum, A. thaliana* (At), *E. coli* (Ec), *Streptococcus equisimilis* (Relseq) RSH. The alignment was performed using the Muscle algorithm (https://www.ebi.ac.uk/Tools/msa/muscle/) with default parameters. Pink horizontal bars indicate important catalytic motifs for (p)ppGpp synthesis (Steinchen & Bange, 2016). Black, dark grey and light grey backgrounds indicate percentage identity between protein sequences (100, 80, 60%, respectively). (B) The structure of the *B. subtilis* SAS1 active site bound to AMCPP compared to the corresponding domain in modelled structures of RSH from *P. tricornutum*, *A. thaliana* RSH1 and *A. thaliana* RSH3. The SAS1 conserved glycine at position 45 is shown in red, and other residues implicated in AMCPP binding are shown in black. These residues are also indicated by an asterisk in 3A. Non-conserved residues are indicated in single letter code in the modelled RSH active sites. Structures were modelled using the I-Tasser Modelling Suite with standard settings (Yang *et al.*, 2015).

In AtRSH1, which lacks (p)ppGpp synthetase activity (Sugliani *et al.*, 2016), the residue corresponding to R46 is substituted by a valine, whose side-chain does not retain AMCPP and due to its polarity cannot bond with the oxygens in the ribose ring (Fig. 3B). Many other conserved residues involved in AMCPP binding are also missing from AtRSH1. In contrast, the synthetase active site of PtRSH1 more closely resembles that of an active (p)ppGpp synthetase: there is a glycine residue at the position corresponding to SAS1 G45 and the neighbouring arginine involved in AMCPP binding takes a similar orientation to the corresponding arginine in AtRSH3, which acts as a ppGpp synthetase (Fig. 3B). These modelling results strongly support the results of our activity assays (Fig. 2), and show that the residue corresponding to SAS1 G45 can be substituted by at least alanine (PtRSH4a) or serine (PtRSH4b) residues, without necessarily affecting the conformation of the active site or the orientation of the neighbouring arginine, R46.

### Evolution and domain organization of *P. tricornutum* and other algal RSH enzymes

Our work shows that the *P. tricornutum* RSH enzymes have significant differences with known plant and algal RSHs. However, the diatom RSH enzymes have not yet been placed within a phylogenetic context. This is necessary for determining whether they belong to new RSH clades, and for understanding when the RSH enzymes were acquired during the complex evolutionary history of the diatom plastid. Therefore, we constructed a phylogenetic tree based on the sequences of the hydrolase and synthetase domains of different RSH enzymes from bacteria, plants and algae (Fig. 4).

**Fig. 4.**
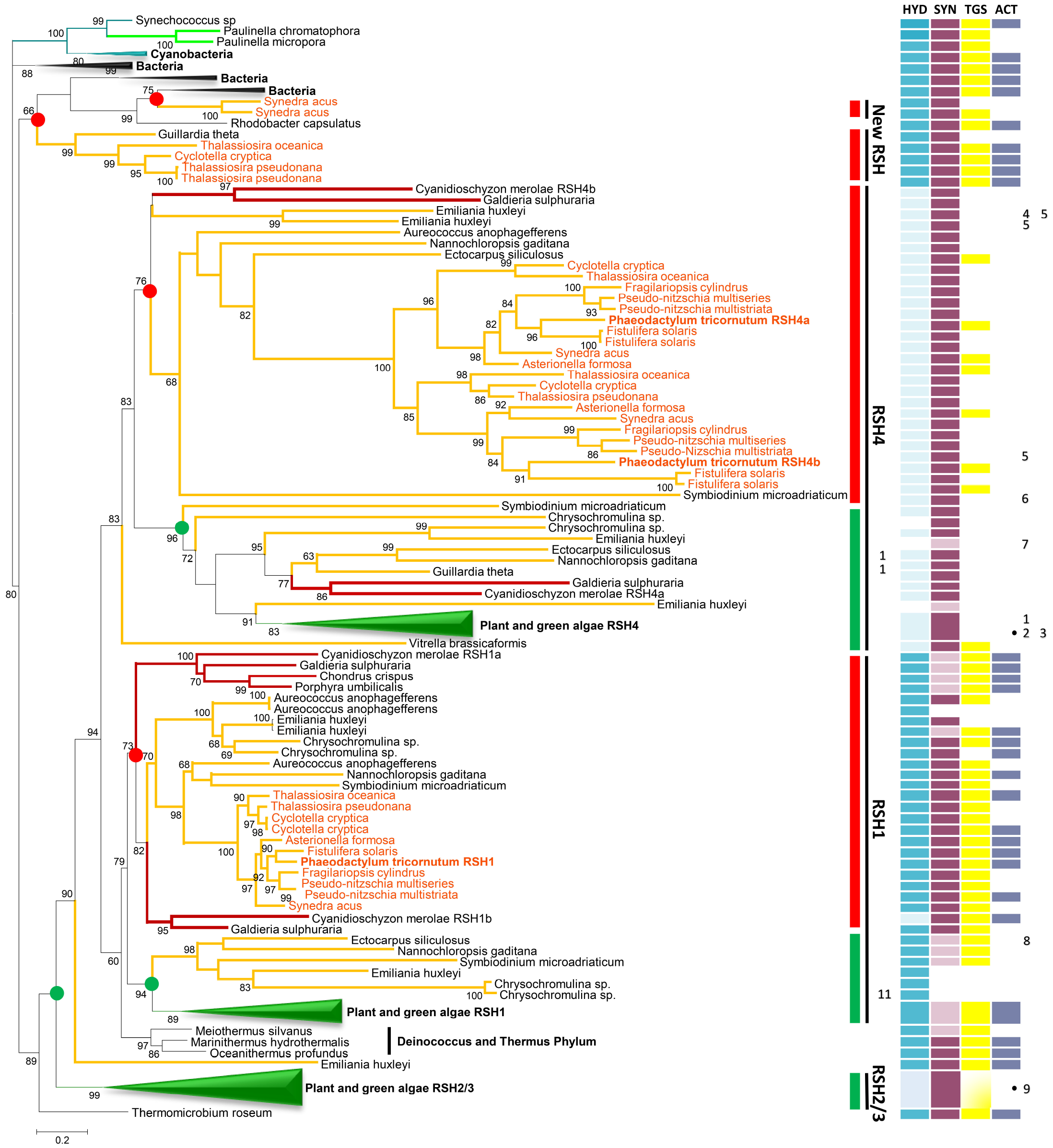
Phylogenetic tree of RSH proteins from diatoms and other photosynthetic organisms. The synthetase and hydrolase domains of different RSH proteins were used to infer phylogenetic relationships using maximum likelihood reconstructions. Diatom RSH are indicated by orange text and branches, CASH RSH are indicated by orange branches, red algal RSH are indicated by red branches, RSH from plants and green algae are condensed and indicated by green branches, cyanobacteria are condensed and indicated by cyan branches, and bacteria are condensed and indicated by black branches. The scale bar indicates the number of substitutions per sites. Statistical support for branches was estimated with the SH-like local support method. Only support values greater than 50% are shown. RSH groups are indicated to the right, and red or green circles at the nodes indicate the base of each major RSH clade Accession numbers and the alignment are provided in supplementary Table S3 and Supporting Dataset 1. Domain architecture is shown to the right of the phylogenetic tree. Domains were identified using the NCBI Conserved Domain (CD) algorithm. Domains: (p)ppGpp hydrolase (HYD), (p)ppGpp synthetase (SYN), Threonyl-tRNA synthetase GTPase Spot (TGS) and Aspartate kinase, Chorismate mutase, TyrA (ACT). Pale coloured HYD and SYN domains columns indicate a likely loss of activity due to the substitution of important residues. Other domains are indicated by numbers and listed in supplementary Table S4. A dot indicates that the domain is present in several members of a condensed clade.

PtRSH4a and PtRSH4b belong to a clade that is monophyletic with a sister clade containing the plant and green algae RSH4 enzymes, and which together can be considered as part of an extended RSH4 clade (Fig. 4). PtRSH4a and PtRSH4b are members of two well separated diatom RSH clades that form part of a larger clade composed of RSH from red algae and the CASH lineages (organisms descended from an endosymbiosis between a eukaryote and a red alga: Cryptomonads, Alveolates, Stramenopiles and Haptophytes)(Petersen *et al.*, 2014). We will refer to this clade as the red RSH4 clade, based on the principal source of chloroplast ancestry. Within the red RSH4 clade the substitution of the residue equivalent to G240 that we observed in PtRSH4a and PtRSH4b is widespread among the CASH lineages but not red algae. Interestingly, the plant and algal sister clade also contains RSH4 enzymes from some red algae and CASH lineages, but not from diatoms. This clade can therefore be considered a red and green RSH4 clade based on chloroplast ancestry. The existence of two monophyletic RSH4 clades indicates that the common ancestor of was likely to have had two RSH4 like enzymes that were subsequently retained, or selectively lost/duplicated in the different lineages.

PtRSH1 likewise belongs to a red algal and CASH lineage clade (or red RSH1 clade) that is monophyletic with a sister clade containing the plant and green algal RSH1 enzymes (red and green RSH1 clade), and which together can be considered as part of an extended RSH1 clade (Fig. 4). The majority of members of the red RSH1 clade possess a synthetase domain where all residues known to be critical for (p)ppGpp synthetase activity are intact, suggesting that all these enzymes have the potential for bifunctionality. The red and green RSH1 clade contains RSH1 enzymes from several CASH organisms, but there are no red algal RSH1 at its base. For this reason, it is more difficult to infer the evolutionary relationship between the two major RSH1 clades. Phylogenetic analysis of full-length RSH1-like sequences complete with the C-terminal regulatory domain also supported the existence of a red RSH1 clade that is distinct from the red and green RSH1 clade (Fig. S3). However, the full-length RSH1 tree also did not clearly resolve the evolutionary history of the extended RSH1 clade. As observed in previous analyses of plant and algal RSH enzymes we found that the RSH1 enzymes group closely with RSH enzymes from the Deinococcus–Thermus bacterial phylum (Givens *et al.*, 2004; Atkinson *et al.*, 2011; Ito *et al.*, 2017).

Our analysis also reveals that other diatom species possess additional RSH enzymes to those found in *P. tricornutum*. Unlike the PtRSH, these additional RSH enzymes are not in clades that are monophyletic with RSH enzymes from plants or green algae, and form two distinct clades within the proteobacterial RSH (top of Fig. 4). These novel RSH clades are unlikely to be sequencing artefacts because there are multiple representatives from different species. Interestingly, we also found no evidence that CASH lineage organisms possess members of the RSH2/3 family suggesting that this family is unique to plants and green algae. Two CASH organisms, *Emiliania huxleyi* and *Vitrella brassicaformis*, also have orphan RSH that are not associated with other algal RSH or with the major RSH families of plants, algae or bacteria (Fig. 4).

Recent studies strongly suggest that plastid arose from the endosymbiosis of a freshwater Gloeomargarita-like cyanobacterium (Ponce-Toledo *et al.*, 2017; Sánchez-Baracaldo *et al.*, 2017). Strikingly, our phylogenetic analysis gives no indication that any of the major RSH groups (RSH1, RSH2/3 and RSH4) are more closely related to Gloeomargarita, or indeed any cyanobacterial species, than the RSH from other bacteria.

### Domain organization of *P. tricornutum* and other algal RSH enzymes

In terms of domain structure, the majority of plant and green algal RSH4 have acquired a C-terminal EF-hand domain (Fig. 4). Two stramenopile RSH4 in the same clade as the plant and algal RSH4 also possess EF-hand domains, though at the N-terminus. No EF-hand domains could be detected in PtRSH4a and PtRSH4b or indeed any members of the sister RSH4 clade that contains only CASH and red algal RSH. However, a C-terminal MoaD-like protein domain was encountered in three separate members of this clade. We also observed isolated domain acquisitions in both clades of the extended RSH4 family: a major intrinsic protein domain in the alveolate *Symbiodinium microadriaticum*, a minor capsid protein VI domain in the alveolate *Vitrella brassicaformis*, a phosphopantetheinyl transferase domain in the haptophyte *Emiliania huxleyi*, and a forkhead domain in the green alga *Chlorella variabilis*. Interestingly, Ito et al. (2017) previously reported the acquisition of tetratricopeptide repeat domains at the C-terminus of RSH4 from some green algae species, acquisitions that were also detected here.

Less evidence of domain acquisition could be observed within the extended RSH1 family, where the majority of enzymes, including PtRSH1, contain both TGS and ACT domains. Nevertheless, an RSH1 from the brown alga *E. siliculosus* contains a Chlorophyll A-B binding protein domain, and an RSH1 from the haptophyte *Chrysochromulina* sp contains an endomucin domain.

## Discussion

In most bacteria, nutrient limitation or stress provokes the rapid synthesis of the two nucleotide alarmones, pppGpp and ppGpp. In *E. coli*, (p)ppGpp synthesis is controlled by two enzymes, RelA and SpoT. Chloroplast localised RelA SpoT homologue (RSH) enzymes are also found in algae and plants, where they are implicated in the control of chloroplast function and plant development (Boniecka *et al.*, 2017; Field, 2018). Little is known about these RSH enzymes in diatoms, or indeed the entire red algal lineage. In this report, we showed that the genome of the diatom *P. tricornutum* encodes three RSH enzymes: PtRSH1, PtRSH4a and PtRSH4b. Analysis of their amino acids sequence revealed the presence of probable chloroplast targeting peptides, indicating that these enzymes are likely to be located within the chloroplast, the site of (p)ppGpp synthesis and its action in plants (Field, 2018) (Fig. 1A). Using heterologous expression in *E. coli* (p)ppGpp mutants we showed that all three *P. tricornutum* RSH are (p)ppGpp synthetases, and that PtRSH1 is also a (p)ppGpp hydrolase (Fig. 2). Gene expression data indicates that the *P. tricornutum RSH* are expressed, and that PtRSH4a is strongly induced in response to certain stresses (Table S2) (Scala *et al.*, 2002; Bowler *et al.*, 2008; Matthijs *et al.*, 2016; Matthijs *et al.*, 2017). Finally, phylogenetic analysis indicated that PtRSH1, PtRSH4a and PtRSH4b are typical members of well-defined diatom RSH clades that belong to the previously identified plant and algal RSH1 and RSH4 families (Fig. 4)(Atkinson *et al.*, 2011; Ito *et al.*, 2017). These results indicate that the *P. tricornutum* RSH enzymes and their orthologues are sufficient for the establishment of a (p)ppGpp homeostasis mechanism in the chloroplasts of diatoms.

PtRSH1 is unusual in that it functions as a bifunctional (p)ppGpp synthetase / hydrolase RSH. Bifunctional RSH such as SpoT are common in bacteria, but no bifunctional RSH enzymes have been demonstrated in the photosynthetic eukaryotes before now. We note however that members of the plant and green algal RSH2/3 family often possess almost intact hydrolase domains in addition to active synthetase domains (Fig. 4). However, hydrolase activity has not so far been demonstrated for one of these enzymes and their overexpression in plants results in (p)ppGpp accumulation (Maekawa *et al.*, 2015; Sugliani *et al.*, 2016). Interestingly, our phylogenetic analysis suggests that all members of the red RSH1 clade may be bifunctional (Fig. 4). This is in contrast to the sister red and green RSH1 clade where the majority of RSH1 appear to have lost (p)ppGpp synthetase activity due to the loss of essential residues in the active site (Sugliani *et al.*, 2016; Ito *et al.*, 2017). Our results are also supported by modelling of the active sites of PtRSH1 and Arabidopsis RSH1 (Fig. 3b). Arabidopsis RSH1, while preserving the global topology of the synthetase active site, has lost the arginine residue with a guanidinium group that is likely to retain ATP in its binding pocket. Structural data indicates that bifunctional RSH can switch between (p)ppGpp-hydrolase-OFF / (p)ppGpp-synthase-ON and hydrolase-ON / synthase-OFF configurations (Hogg *et al.*, 2004). Our heterologous expression results suggest that PtRSH1 is likely to be capable of undergoing similar activity switches (Fig. 2). The regulation of PtRSH1 and other members of the red RSH1 clade may therefore be significantly different to plant and green algal RSH1 and may reflect the different lifestyles of these organisms.

PtRSH4a and PtRSH4b are also unusual among (p)ppGpp synthetases due to the substitution of a residue corresponding to SAS1 G45 / Rel_seq_ G240. This conserved residue has previously been shown to be necessary for (p)ppGpp synthetase activity (Wendrich & Marahiel, 1997; Hogg *et al.*, 2004) and substitution of this residue is widely used to infer the loss of (p)ppGpp synthetase activity (Masuda *et al.*, 2008; Mizusawa *et al.*, 2008; Sato *et al.*, 2015; Ito *et al.*, 2017; Imamura *et al.*, 2018). Modelling of the active site of PtRSH4a and PtRSH4b shows that the glycine substitution does not appear to affect ATP binding by the neighbouring arginine (Fig. 3b), and sequence analysis indicates that the glycine substitution is widespread in the red RSH4 clade. These data suggest that the previous reports of an association between the substitution of the residue corresponding to SAS1 G45 and the loss of synthetase activity may be conditional on the presence of other specific residues in the active site. PtRSH4a and PtRSH4b, and nearly all members of red RSH4 clade lack the EF-hand domain found at the C-terminal of the majority of members of the sister red and green RSH4 clade. A certain number of novel domain acquisitions were also observed within both the major RSH4 clades in addition to those previously identified in the red and green RSH4 clade (Ito *et al.*, 2017). Altogether, these findings suggest that extended RSH4 family enzymes act exclusively as (p)ppGpp synthetases, and are susceptible to domain acquisition, presumably for new regulatory functions. Indeed, altered regulation might be expected, because many enzymes from diatoms involved in processes such as CO_2_ assimilation, sulphate assimilation have different regulatory properties than their orthologues in plants (Maberly *et al.*, 2010; Jensen *et al.*, 2017). Determining the functions of these domains in (p)ppGpp metabolism presents a fascinating challenge for future research.

We show here that, when the RSH of genome-sequenced diatoms are considered, the evolutionary history of RSH enzymes in the photosynthetic eukaryotes is considerably more complex than previously thought (Givens *et al.*, 2004; Atkinson *et al.*, 2011; Ito *et al.*, 2017). We extend the described RSH1 and RSH4 families by showing the existence of sister clades specific to the red-lineage that show distinct functional and structural properties (Fig. 2-4). Interestingly, although Atkinson et al. (2011) included several CASH and red algal RSH1 enzymes in their phylogenetic analysis, they did not detect a clear separation of the RSH1 family into a red RSH1 and a red and green RSH1 clade, or the existence of a red RSH4 clade. The difference between these studies is very likely to be due to our use of many more CASH RSH sequences, which are available today thanks to the ever growing list of sequenced CASH genomes. For the same reason we also detected two new CASH RSH families that are not monophyletic with any of the known RSH families in photosynthetic eukaryotes.

Previous reports on the phylogenetic relationships of bacterial, plant and algal RSH enzymes have proposed that plant and algal RSH may have arisen through lateral gene transfer rather than vertical descent from the ancestral chloroplast (Givens *et al.*, 2004; Atkinson *et al.*, 2011; Ito *et al.*, 2017). A major reason for this proposition is the grouping of the RSH1 family with RSH from the prokaryotic Deinococcus–Thermus phylum rather than cyanobacteria. Our analysis supports this idea by also showing that the extended RSH1 family groups with RSH from the Deinococcus-Thermus phylum (Fig. 4). Furthermore, our discovery of new CASH RSH families that group with bacteria are evidence that additional lateral gene transfers from bacteria may have occurred recently. This would suggest that lateral transfers of RSH genes can occur readily.

Our report sheds new light on the RSH enzyme family and (p)ppGpp metabolism in the diatoms, and reveals many surprises. Further research is now required to elucidate the mechanism and role of (p)ppGpp signalling in the lifestyle of this important and diverse group of photosynthetic eukaryotes.

## Supporting information

## Acknowledgements

This work was supported by the Agence Nationale de la Recherche (ANR-15-CE05-0021-03).

## Author Contributions

A. L, B.M, A.V, E.B, B.F and B.G conceived and planned the experiments. A.L, C.P, A.V and B. F performed the experiments. A.L, C.P, A.V, E.B, B.F and B.G contributed to the interpretation of the results. B.F, A.L and B.G wrote the manuscript. All authors provided critical feedback and helped shape the research, analysis and manuscript.

## Supplementary data

**Fig. S1** Full amino acid sequences of the three RSH proteins found in *P. tricornutum*.

**Fig. S2** Alignment of P. tricornutum RSH proteins.

**Fig. S3** Phylogenetic tree of RSH1 proteins from different diatoms and other photosynthetic eukaryotes.

**Table S1** List of the primers used in this study.

**Table S2** RSH EST data.

**Table S3** Accession numbers for sequences used in Fig. 4.

**Table S4** Domains found in RSH enzymes.

**Supporting Dataset 1** Alignment of RSH enzymes sequences (hydrolase and synthetase domains) in fasta format.

**Supporting Dataset 2** Alignment of RSH1 enzymes (whole sequence) in fasta format.

## Abbreviations

ACT: Aspartate kinase, Chorismate mutase, TyrA domain
CASH: Cryptomonads, Alveolates, Stramenopiles and Haptophytes
CRSH: Ca^2+^ activated RelA/SpoT homolog
EF-hand: Ca^2+^ binding domain
(p)ppGpp: guanosine tetraphosphate and pentaphosphate
Rel_seq_: *Streptococcus equisimilis* Rel
RSH: RelA/SpoT homologue
TGS: Threonyl-tRNA synthetase, GTPase, SpoT and RelA domain

## References

Abdelkefi H, Sugliani M, Ke H, Harchouni S, Soubigou-Taconnat L, Citerne S, Mouille G, Fakhfakh H, Robaglia C, Field B. 2018. Guanosine tetraphosphate modulates salicylic acid signalling and the resistance of Arabidopsis thaliana to Turnip mosaic virus. Molecular Plant Pathology 19, 634–646.

Archibald JM. 2015. Genomic perspectives on the birth and spread of plastids. Proceedings of the National Academy of Sciences 112, 10147–10153.

Atkinson GC, Tenson T, Hauryliuk V. 2011. The RelA/SpoT homolog (RSH) superfamily: distribution and functional evolution of ppGpp synthetases and hydrolases across the tree of life. PLoS One 6, e23479.

Battesti A, Bouveret E. 2006. Acyl carrier protein/SpoT interaction, the switch linking SpoT-dependent stress response to fatty acid metabolism. Mol Microbiol 62, 1048–1063.

Boniecka J, Prusinska J, Dabrowska GB, Goc A. 2017. Within and beyond the stringent response-RSH and (p)ppGpp in plants. Planta 246, 817–842.

Bowler C, Allen AE, Badger JH, Grimwood J, Jabbari K, Kuo A, Maheswari U, Martens C, Maumus F, Otillar RP, et al. 2008. The Phaeodactylum genome reveals the evolutionary history of diatom genomes. Nature 456, 239.

Cashel M, Gallant J. 1969. Two compounds implicated in the function of the RC gene of Escherichia coli. Nature 221, 838.

Dalebroux ZD, Swanson MS. 2012. ppGpp: magic beyond RNA polymerase. Nat Rev Microbiol 10, 203–212.

Dorrell RG, Gile G, McCallum G, Meheust R, Bapteste EP, Klinger CM, Brillet-Guéguen L, Freeman KD, Richter DJ, Bowler C. 2017. Chimeric origins of ochrophytes and haptophytes revealed through an ancient plastid proteome. Elife 6, e23717.

Dorrell RG, Smith AG. 2011. Do red and green make brown?: perspectives on plastid acquisitions within chromalveolates. Eukaryot Cell 10, 856–868.

Field B. 2018. Green magic: regulation of the chloroplast stress response by (p)ppGpp in plants and algae. J Exp Bot 69, 2797–2807.

Givens RM, Lin MH, Taylor DJ, Mechold U, Berry JO, Hernandez VJ. 2004. Inducible expression, enzymatic activity, and origin of higher plant homologues of bacterial RelA/SpoT stress proteins in Nicotiana tabacum. J Biol Chem 279, 7495–7504.

Gruber A, Rocap G, Kroth PG, Armbrust EV, Mock T. 2015. Plastid proteome prediction for diatoms and other algae with secondary plastids of the red lineage. Plant J 81, 519–528.

Hauryliuk V, Atkinson GC, Murakami KS, Tenson T, Gerdes K. 2015. Recent functional insights into the role of (p)ppGpp in bacterial physiology. Nat Rev Microbiol 13, 298–309.

Hogg T, Mechold U, Malke H, Cashel M, Hilgenfeld R. 2004. Conformational antagonism between opposing active sites in a bifunctional RelA/SpoT homolog modulates (p)ppGpp metabolism during the stringent response [corrected]. Cell 117, 57–68.

Huesgen PF, Alami M, Lange PF, Foster U, Schroder WP, Overall CM, Green BR. 2013. Proteomic amino-termini profiling reveals targeting information for protein import into complex plastids. PLoS One 8, e74483.

Ihara Y, Ohta H, Masuda S. 2015. A highly sensitive quantification method for the accumulation of alarmone ppGpp in Arabidopsis thaliana using UPLC-ESI-qMS/MS. J Plant Res 128, 511–518.

Imamura S, Nomura Y, Takemura T, Pancha I, Taki K, Toguchi K, Tozawa Y, Tanaka K. 2018. The checkpoint kinase TOR (target of rapamycin) regulates expression of a nuclear-encoded chloroplast RelA-SpoT homolog (RSH) and modulates chloroplast ribosomal RNA synthesis in a unicellular red alga. The Plant Journal 94, 327–339.

Ito D, Ihara Y, Nishihara H, Masuda S. 2017. Phylogenetic analysis of proteins involved in the stringent response in plant cells. J Plant Res 130, 625–634.

Jensen E, Clément R, Maberly SC, Gontero B. 2017. Regulation of the Calvin-Benson-Bassham cycle in the enigmatic diatoms: biochemical and evolutionary variations on an original theme. Philosophical Transactions of the Royal Society B: Biological Sciences 372.

Jeong JY, Yim HS, Ryu JY, Lee HS, Lee JH, Seen DS, Kang SG. 2012. One-step sequence- and ligation-independent cloning as a rapid and versatile cloning method for functional genomics studies. Appl Environ Microbiol 78, 5440–5443.

Katoh K, Rozewicki J, Yamada KD. 2017. MAFFT online service: multiple sequence alignment, interactive sequence choice and visualization. Brief Bioinform. DOI: 10.1093/bib/bbx108

Kim T-H, Ok SH, Kim D, Suh S-C, Byun MO, Shin JS. 2009. Molecular characterization of a biotic and abiotic stress resistance-related gene RelA/SpoT homologue (PepRSH) from pepper. Plant Science 176, 635–642.

Kojadinovic-Sirinelli M, Villain A, Puppo C, Fon Sing S, Prioretti L, Hubert P, Grégori G, Zhang Y, Jean-Françis S, Jean-Michel C, et al. 2018. Exploring the microbiome of the “star” freshwater diatom Asterionella formosa in a laboratory context Enviromental Microbiology in press.

Maberly SC, Courcelle C, Groben R, Gontero B. 2010. Phylogenetically-based variation in the regulation of the Calvin cycle enzymes, phosphoribulokinase and glyceraldehyde-3-phosphate dehydrogenase, in algae. J Exp Bot 61, 735–745.

Maekawa M, Honoki R, Ihara Y, Sato R, Oikawa A, Kanno Y, Ohta H, Seo M, Saito K, Masuda S. 2015. Impact of the plastidial stringent response in plant growth and stress responses. Nature Plants 1, 15167.

Malviya S, Scalco E, Audic S, Vincent F, Veluchamy A, Poulain J, Wincker P, Iudicone D, de Vargas C, Bittner L, et al. 2016. Insights into global diatom distribution and diversity in the world’s ocean. Proc Natl Acad Sci USA 113, E1516–E1525.

Marchler-Bauer A, Derbyshire MK, Gonzales NR, Lu S, Chitsaz F, Geer LY, Geer RC, He J, Gwadz M, Hurwitz DI, et al. 2015. CDD: NCBI’s conserved domain database. Nucleic Acids Res 43, D222–226.

Masuda S, Mizusawa K, Narisawa T, Tozawa Y, Ohta H, Takamiya K. 2008. The bacterial stringent response, conserved in chloroplasts, controls plant fertilization. Plant Cell Physiol 49, 135–141.

Matthijs M, Fabris M, Broos S, Vyverman W, Goossens A. 2016. Profiling of the early nitrogen stress response in the diatom Phaeodactylum tricornutum reveals a novel family of RING-domain transcription factors. Plant Physiology 170, 489–498.

Matthijs M, Fabris M, Obata T, Foubert I, Franco-Zorrilla JM, Solano R, Fernie AR, Vyverman W, Goossens A. 2017. The transcription factor bZIP14 regulates the TCA cycle in the diatom Phaeodactylum tricornutum. EMBO J 36, 1559–1576

Mizusawa K, Masuda S, Ohta H. 2008. Expression profiling of four RelA/SpoT-like proteins, homologues of bacterial stringent factors, in Arabidopsis thaliana. Planta 228, 553–562.

My L, Rekoske B, Lemke JJ, Viala JP, Gourse RL, Bouveret E. 2013. Transcription of the Escherichia coli fatty acid synthesis operon fabHDG is directly activated by FadR and inhibited by ppGpp. J Bacteriol 195, 3784–3795.

Petersen J, Ludewig A-K, Michael V, Bunk B, Jarek M, Baurain D, Brinkmann H. 2014. Chromera velia, Endosymbioses and the Rhodoplex Hypothesis—Plastid Evolution in Cryptophytes, Alveolates, Stramenopiles, and Haptophytes (CASH Lineages). Genome Biol Evol 6, 666–684.

Ponce-Toledo RI, Deschamps P, Lopez-Garda P, Zivanovic Y, Benzerara K, Moreira D. 2017. An Early-Branching Freshwater Cyanobacterium at the Origin of Plastids. Current Biology 27, 386–391.

Price MN, Dehal PS, Arkin AP. 2010. FastTree 2––approximately maximum-likelihood trees for large alignments. PLoS One 5, e9490.

Puppo C, Voisin T, Gontero B. 2017. Genomic DNA extraction from the pennate diatom Asterionella formosa optimised for next generation sequencing, protocols.io. DOI: 10.17504/protocols.io.jytcpwn

Puthiyaveetil S, Kavanagh TA, Cain P, Sullivan JA, Newell CA, Gray JC, Robinson C, van der Giezen M, Rogers MB, Allen JF. 2008. The ancestral symbiont sensor kinase CSK links photosynthesis with gene expression in chloroplasts. Proc. Natl. Acad. Sci. USA 105, 10061–10066.

Ronneau S, Caballero-Montes J, Mayard A, Garcia-Pino A, Hallez R. 2018. Regulation of (p)ppGpp hydrolysis by a conserved archetypal regulatory domain. bioRxiv. DOI: 10.1101/257592

Sánchez-Baracaldo P, Raven JA, Pisani D, Knoll AH. 2017. Early photosynthetic eukaryotes inhabited low-salinity habitats. Proceedings of the National Academy of Sciences 114, E7737–E7745.

Sato M, Takahashi T, Ochi K, Matsuura H, Nabeta K, Takahashi K. 2015. Overexpression of RelA/SpoT homologs, PpRSH2a and PpRSH2b, induces the growth suppression of the moss Physcomitrella patens. Biosci Biotechnol Biochem 79, 36–44.

Scala S, Carels N, Falciatore A, Chiusano ML, Bowler C. 2002. Genome Properties of the Diatom Phaeodactylum tricornutum. Plant Physiology 129, 993–1002.

Steinchen W, Bange G. 2016. The magic dance of the alarmones (p)ppGpp. Mol Microbiol 101, 531–544.

Steinchen W, Schuhmacher JS, Altegoer F, Fage CD, Srinivasan V, Linne U, Marahiel MA, Bange G. 2015. Catalytic mechanism and allosteric regulation of an oligomeric (p)ppGpp synthetase by an alarmone. Proc Natl Acad Sci U S A 112, 13348–13353.

Sugliani M, Abdelkefi H, Ke H, Bouveret E, Robaglia C, Caffarri S, Field B. 2016. An Ancient Bacterial Signaling Pathway Regulates Chloroplast Function to Influence Growth and Development in Arabidopsis. Plant Cell 28, 661–679.

Takahashi K, Kasai K, Ochi K. 2004. Identification of the bacterial alarmone guanosine 5′-diphosphate 3′-diphosphate (ppGpp) in plants. Proc. Natl. Acad. Sci. USA 101, 4320–4324.

van der Biezen EA, Sun J, Coleman MJ, Bibb MJ, Jones JD. 2000. Arabidopsis RelA/SpoT homologs implicate (p)ppGpp in plant signaling. Proc. Natl. Acad. Sci. USA 97, 3747–3752.

Vardi A, Formiggini F, Casotti R, De Martino A, Ribalet F, Miralto A, Bowler C. 2006. A stress surveillance system based on calcium and nitric oxide in marine diatoms. PLoS Biol 4, e60.

Wahl A, My L, Dumoulin R, Sturgis JN, Bouveret E. 2011. Antagonistic regulation of dgkA and plsB genes of phospholipid synthesis by multiple stress responses in Escherichia coli. Mol Microbiol 80, 1260–1275.

Wendrich TM, Marahiel MA. 1997. Cloning and characterization of a relA/spoT homologue from Bacillus subtilis. Mol Microbiol 26, 65–79.

Wout P, Pu K, Sullivan SM, Reese V, Zhou S, Lin B, Maddock JR. 2004. The Escherichia coli GTPase CgtAE cofractionates with the 50S ribosomal subunit and interacts with SpoT, a ppGpp synthetase/hydrolase. J Bacteriol 186, 5249–5257.

Yamburenko MV, Zubo YO, Borner T. 2015. Abscisic acid affects transcription of chloroplast genes via protein phosphatase 2C-dependent activation of nuclear genes: repression by guanosine-3′-5′-bisdiphosphate and activation by sigma factor 5. Plant J 82,1030–1041.

Yang J, Yan R, Roy A, Xu D, Poisson J, Zhang Y. 2015. The I-TASSER Suite: protein structure and function prediction. Nat Methods 12, 7–8.

