## Supplementary material for "Golden magic: RSH enzymes for (p)ppGpp metabolism in the diatom *Phaeodactylum tricornutum*"

**Supplementary Data**

### Fig. S1 Full amino acid sequences of the three RSH proteins found in P. tricornutum.

```
>PtRSH1-11099
MQVSSTGGMWTILLLAFLWSCTCIAFTTPRWFPTPNMRHSQSIVPLFDRIPPSFPSLEMAGANTTGSGMNSAQTLFP
TPDRTSLASIYTSESHTAAAAVMSNGFNYSAHYNTAALIPPVTFTSASTEVAAEKDRSEADTPPEVNHWTEIMRVGW
KDDTSTTNSSTTSAQISSSSTKWDAVWSDYDLQQIERYWDRIMPTVSYLGTDAVAKIYQALCVAYRAHRGQMRKSGE
PFIVHPVEVSLLLSGLKMDAETVMSGLLHDTVEDTDLTFQQVETLFGHTVRSIVEGETKVSKLPKLAFAEYADEQAE
NLRQMFVAMTSDYRIIIVKLADRLHNMRTLRYMKSEKQIKISRETLDIFAPLAHRMGIWQFKSELEDTSFMYLYPQE
YKRLNRRLRLHQQSFQETLEKAQDILQRQLNLDSTLQQQAYKVEVSGRTKEIYSLWHKMETKNVQNLDHIVDVVALR
VVISPRDKSVEADKSELSNDSDRGVWLCYHALGLVQHLPGFVPVPTRVKDYISFPKPNGYQSLHTALILNGQTIEVQ
IRTSMMHQVAEYGMASHWAYTDDKRRGSNEELYSTPWLSSIKEWQNEAVSSRDFVDSVRRELLGKRVFVFLRNGKIL
NLARGATAIDAAFQIHTEVGLSMHGVEINGKPVPLSYELQNGDVVSILTGSGRPATDWMRYAKSRSTRSKLRSYFRD
RQKESLREAGKILLMDYLTVHGTLIQESSYLEKDFAIPKSTEELEYLLPGKTSFNDVDELLVGIGKNHNRSKLHQIV
SQLFEVPKRILITAEEKKIPRLPSNIFAAVQLRQDRAKDAGDAVDLVGEIVTDPATWSTLPKKSVFSYSAESMLAGL
DLPIEYADPEHLCVDCLPVYEDEIVGTRKSGSVSIPMVHRVGCPHAQRAINQAKAHQRQKPFVSKLQNVTSVPGTSI
PRPQLRVDSVSLRQTYGKTAPWMRRAGGQTSKYKSGTVDLPVKLQWSDLDEKDSLFLSEIVVHCGDRKLLLADCSEV
VSETVEIIKTGSSTNEETATLVFLVRVGGLGHIQTLMDRLMKVRSVLSVERRFGSELR

>PtRSH4a-7629
MTNGIPAATATAAAATPTLPLSTHRRRRRMPTVLSTAALLLSSRCSQSADAFTALHNGFRSSKTSLAFRSNVMDGLR
TDALLSTSSSSPTSTIGPLPTWLSYPQAHLKDSLVAELSQAMKVSFFTETETLQLLAAVEEAAGGDAHKVAGTADFL
RILVETMEMGLNALVAAAFHYADCVELREHTRLQSSTSQTAAAMVRHANLDAYYGEHVSQIADDAGRLKQLEWVAQV
VMQTHASRASPDAHDAENLRQLLLSETRDWRALAIRAGACLYRLRGLLKSDSYELTPERVRVGREALSIYAPLASRL
GMHRLKNELEGAAFRVLYQRQYQAVNAMAKEVQTKDENNNMRDVLAEVKNDLTLLLQKDPEFSKAVSDFTVTARVKE
SYSMWKKMLRHGYKHLLQVPDALALRIVLNAKKETPNEPVEVTRARERALCYYAQKLCTSQFAPVANAPRFKDYVER
PKPNGYQSLHYTATYESWKVEIQVRSGEMHQVAEFGLASHWDYKASQDSLAEADVEPSDLDQSSDAYVRKVQEWHWD
QHNGVAPATVEWDASPASFAPVTASDIWQSRIRAERIRARTQRLEPYLQALTAAQSDLAREYVFCFLKSGDTPKVLA
LPAGACVLDALRQGGVDGSVQLNGVEASITRQLTNGDVLTISSLAVV

>PtRSH4b-33947
MKSGLLDYTTLTVGISLLHKRNGNTMKAKKRPRAERIQVGKPSASAFSWWSLFLIEWSSLPVFGLTQWSLYGNEFTT
QSKPTELRAIQSPVRNDAVSPRKKKEILLDFPDSPSSRTGEHVNSEFSPQNRIGLPPCCLPYDHFTAEEIEVEVGWL
QYSLLDHGVSFDDVRQIVSTIYNVSENNTSVTVGIVQFLRLLFDTCGEEAHMDRMLSTSVVLASVYHYAECMEAHNQ
GSTAYLLGNANLRNKEMPHNRASIDSEGTVLPAIRGEDTLTRDIISPKPMPRRLRSSTGSFGAGDEVFLITEGAARI
KRAEALVQSVIGNGHIISQAESDLFRDWLLSVMDDWRSLAIRVFACLYRLEGIRLDAGTYDGRTPEVVKLAKEAMRV
YSPLAGRLGMYRLKSRLDEEAFRILYRRQYNAVSSLYLESGAAMEAVSNILRTKISVALQQDESLMMQLEGLEVSSR
VKQPYSFWKKLLKKRTGGLSIVDRRAITNDSTLSIAQVQDGIALRVIIQARKWTENEPLEEIRARERFFCYYVQHQI
RMKWPEVEADRVKDYILYPKPNGYRSLHHTSSVNCNGVDFPFEVQVRSDEMHMIAEYGVAAHWGYKLGNILAPSSPS
VGACAGMLPPAKQCCEILSPAFRPGFSFSHTRLGSTQSFADALVDAKETLLEQYVYVFISGISDESDGQLLSLPAKS
LVVDALAAMDKIDASNLRVLLNGKRVKLDDVVENGDVLMVVA
```

**Fig. S2 Alignment of *P. tricornutum* RSH enzymes.** Amino acid alignments for RSH from *P. tricornutum*, *A. thaliana* (At), *E. coli* (Ec), and *Streptococcus equisimilis* RSH. The alignment was performed using the Muscle algorithm (https://www.ebi.ac.uk/Tools/msa/muscle/ ) with default parameters. Blue and pink horizontal bars indicate important catalytic motifs for (p)ppGpp hydrolysis and synthesis (Steinchen & Bange, 2016). Black, dark grey and light grey backgrounds indicate percentage identity between protein sequences (100, 80, 60%, respectively).

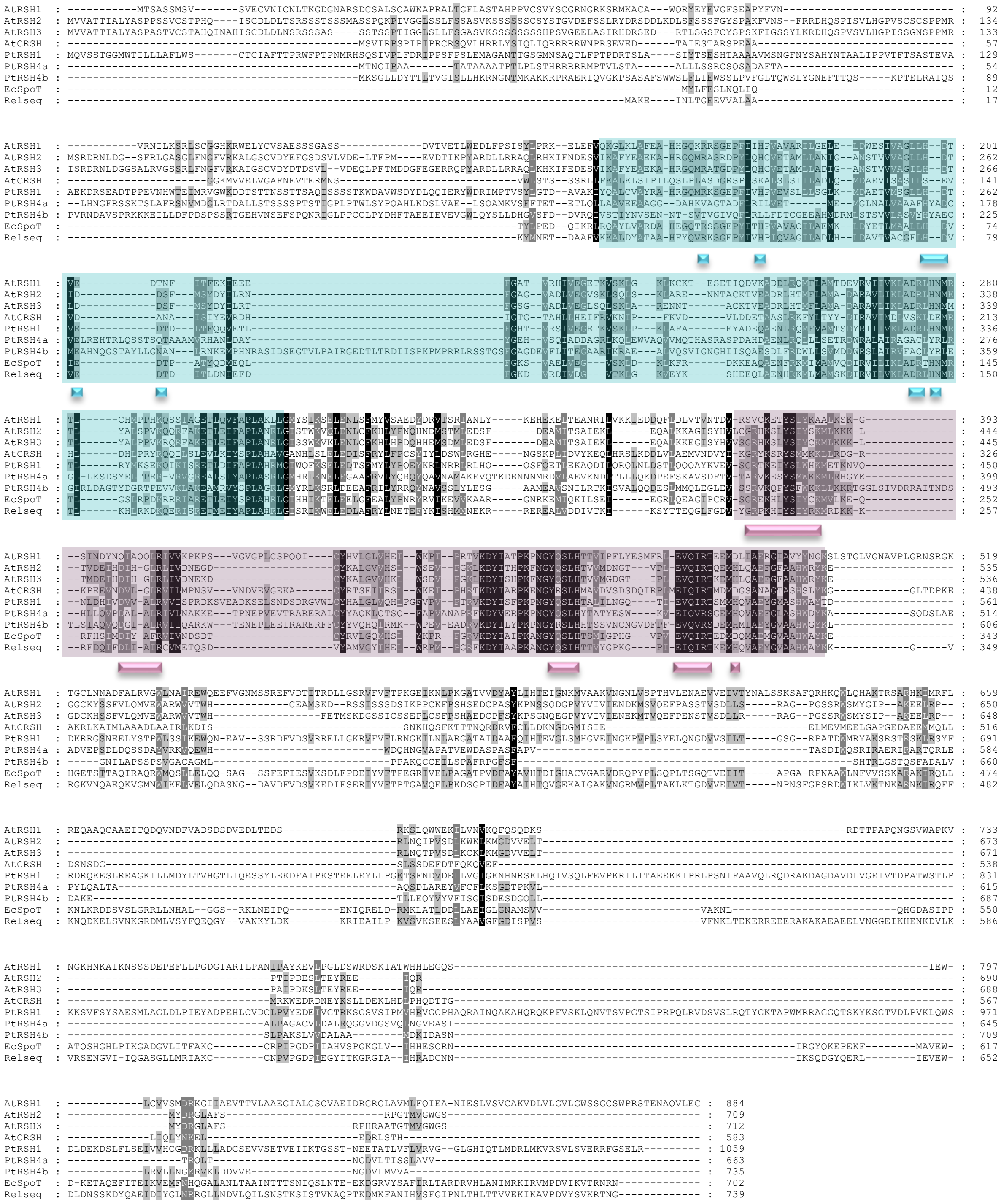

**Fig. S3 Phylogenetic tree of RSH1 proteins from different diatoms and other photosynthetic eukaryotes.** The whole sequences of the different RSH1 proteins were used for the multiple alignment and phylogenetic inference. CASH RSH are indicated by orange branches, diatom RSH are indicated by brown text. The other sequences are from plant and green algae (green branches), red algae (red branches), and bacteria (black branches). The scale bar indicates the number of substitutions per sites. Statistical support for branches was estimated with the SH-like local support method. Only support values greater than 50% are shown. The tree is rooted to *E. coli* SpoT.

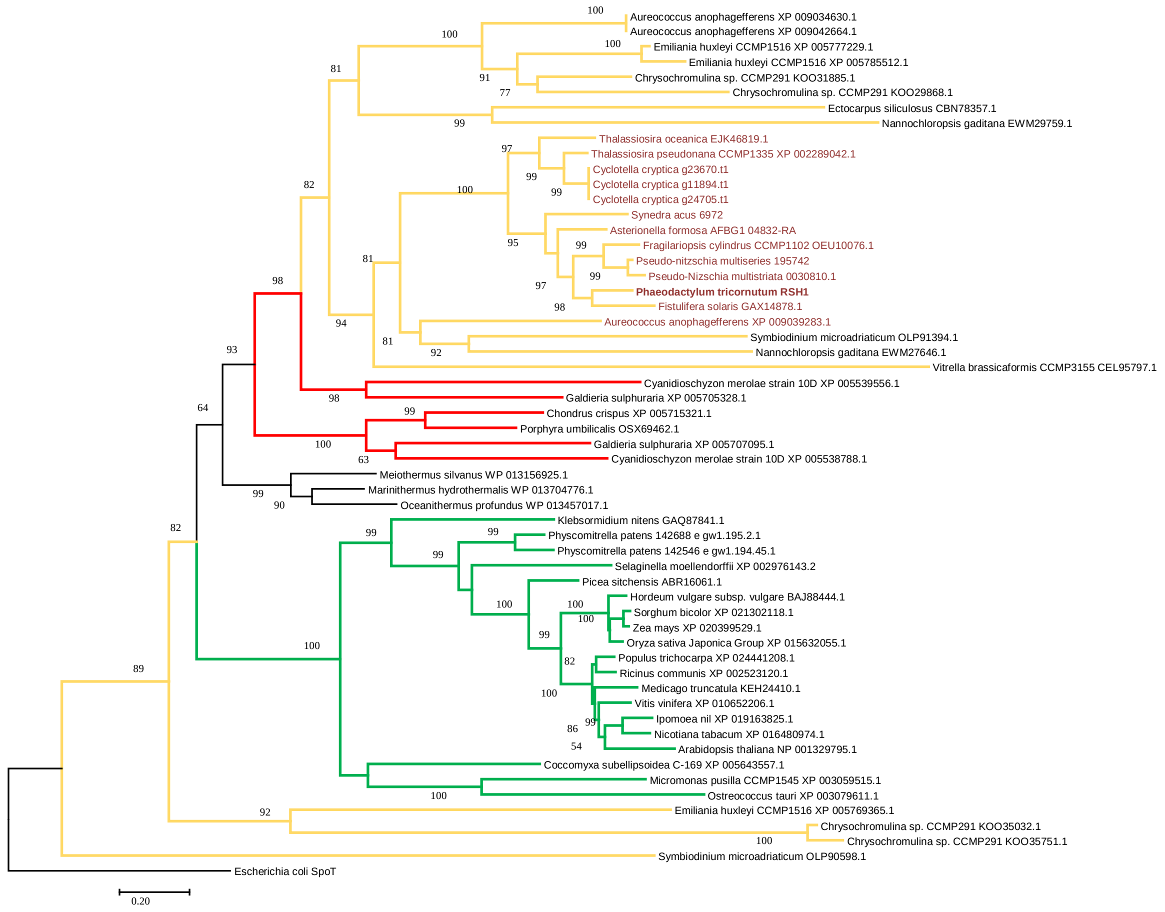

**Table S1 List of the primers used in this study.**

| Name | Sequence (5’ -3’) | Description |
|---|---|---|
| 11099 fw exon1<br>11099 rev exon2 | GCAGGAGGAATTCACCATGGCAATCTACACCTCGGAATCCCA<br>CCGCCAAAACAGCCAAGCTTCTAGCGCAACTCCGAACCAA | Cloning of the PtRSH1(complete gene) |
| 11099 rv exon1<br>11099 fw exon2 | CTTGCCATACCGTATTCAGCTACTTGATGC<br>GCTGAATACGGTATGGCAAGTCACTGGGCG | SLIC for the spliced PtRSH1 gene. |
| 7629 fw<br>7629 rev | AGCAGGAGGAATTCACCATGGACGGTCTCCGTACCGATGC<br>CCGCCAAAACAGCCAAGCTTTCAAACCACCGCGAGGGACG | Cloning of the PtRSH4a |
| 33947 fw<br>33947 rev | CAGGAGGAATTCACCATGGGTCCAGACTCTCCAAGCAGTC<br>CCGCCAAAACAGCCAAGCTTTTATGCTACAACCATCAAAACATC | Cloning of the PtRSH4b |

**Table S2 Number of expressed sequence tags representing the RSH genes per 10,000 total EST in *P. tricornutum* EST databases.**

| ***P tricornutum*** **cDNA library** | **EST Database** | **RSH JGI accession number** | | | **Frustulin 3 JGI** |
|---|---|---|---|---|---|
| | | **7629** | **33947** | **11099** | **51797** |
| Standard condition | 9653 | 7 | 1 | 1 | 61 |
| Iron starved | 21076 | 15 | 0 | 1 | 28 |
| Urea adapted | 21085 | 7 | 0 | 2 | 44 |
| Ammonium adapted | 21081 | 2 | 0 | 1 | 27 |
| Nitrate starved | 21080 | 10 | 0 | 0 | 39 |
| Silica plus | 21077 | 10 | 0 | 1 | 36 |
| Silica minus | 21082 | 13 | 0 | 0 | 38 |
| Exposed to CO2 (24hours) | 23054 | 12 | 0 | 0 | 22 |
| Exposed to CO2 (4 days) | 23055 | 17 | 2 | 3 | 30 |
| Cold stress | 21078 | 2 | 0 | 2 | 54 |
| Low salinity | 21079 | 2 | 0 | 2 | 87 |
| High 2E, 4E decadienal | 21084 | 22 | 0 | 0 | 22 |
| Low 2E, 4E decadienal | 21075 | 1 | 0 | 2 | 39 |
| Triradiate cell-enriched | 23053 | 11 | 0 | 0 | 26 |

Standard conditions: f/2 medium at 20°C 12 h light/12 h dark photoperiod,150 µmol $m^{-2}$ $s^{-1}$ (Scala et al. 2002). Other EST databases from Bowler et al. 2008.

**Table S3 Accession numbers of sequences used for the phylogeny in Fig.4.**

| Species | Taxonomic group | Accession number | RSH type |
|---|---|---|---|
| | **Chromalveolate/ Stramenopile** | | |
| *Asterionella formosa* | Diatom | AFBG1_04832-RA | RSH1 |
| *Asterionella formosa* | Diatom | AFBG1_05489-RA | RSH4 |
| *Asterionella formosa* | Diatom | AFBG1_08512-RA | RSH4 |
| *Cyclotella cryptica* | Diatom | g11894.t1 | RSH1 |
| *Cyclotella cryptica* | Diatom | g23670.t1 | RSH1 |
| *Cyclotella cryptica* | Diatom | g2078.t1 | RSH4 |
| *Cyclotella cryptica* | Diatom | g5699.t1 | RSH4 |
| *Cyclotella_cryptica* | Diatom | g9445.t1 | bacterial |
| *Fistulifera solaris* | Diatom | GAX14878.1 | RSH1 |
| *Fistulifera solaris* | Diatom | GAX10747.1 | RSH4 |
| *Fistulifera solaris* | Diatom | GAX12045.1 | RSH4 |
| *Fistulifera solaris* | Diatom | GAX18927.1 | RSH4 |
| *Fistulifera solaris* | Diatom | GAX23555.1 | RSH4 |
| *Fragilariopsis cylindrus* | Diatom | OEU10076.1 | RSH1 |
| *Fragilariopsis cylindrus* | Diatom | OEU20746.1 | RSH4 |
| *Fragilariopsis cylindrus* | Diatom | OEU21949.1 | RSH4 |
| *Phaeodactylum_tricornutum* | Diatom | JGI:11099 | RSH1 |
| *Phaeodactylum_tricornutum* | Diatom | JGI :7629 | RSH4a |
| *Phaeodactylum_tricornutum* | Diatom | JGI :33947 | RSH4b |
| *Pseudo-nitzschia multiseries* | Diatom | 195742 | RSH1 |
| *Pseudo-nitzschia multiseries* | Diatom | 178926 | RSH4 |
| *Pseudo-nitzschia multiseries* | Diatom | 193757 | RSH4 |
| *Pseudo-nitzschia multistriata* | Diatom | 0030810.1 | RSH1 |
| *Pseudo-nitzschia multistriata* | Diatom | 0019220.1 | RSH4 |
| *Pseudo-nitzschia multistriata* | Diatom | 0074170.1 | RSH4 |
| *Synedra acus* | Diatom | 6972 | RSH1 |
| *Synedra acus* | Diatom | 11925 | RSH4 |
| *Synedra acus* | Diatom | 7671 | RSH4 |
| *Synedra acus* | Diatom | 6453 | bacterial |
| *Synedra acus* | Diatom | 872 | bacterial |
| *Thalassiosira oceanica* | Diatom | EJK46819.1 | RSH1 |
| *Thalassiosira oceanica* | Diatom | EJK45244.1 | RSH4 |
| *Thalassiosira oceanica* | Diatom | EJK49442.1 | RSH4 |
| *Thalassiosira oceanica* | Diatom | EJK63859.1 | bacterial |
| *Thalassiosira pseudonana* | Diatom | XP_002289042.1 | RSH1 |
| *Thalassiosira pseudonana* | Diatom | XP_002293072.1 | RSH4 |
| *Thalassiosira pseudonana* | Diatom | XP_002294153.1 | bacterial |
| *Thalassiosira pseudonana* | Diatom | XP_002297567.1 | bacterial |
| *Ectocarpus siliculosus* | Brown alga | CBN78357.1 | RSH1 |
| *Ectocarpus siliculosus* | Brown alga | CBJ27227.1 | RSH4 |
| *Ectocarpus siliculosus* | Brown alga | CBJ31981.1 | RSH4 |
| *Nannochloropsis gaditana* | Eustigmatophyte | EWM27646.1 | RSH1 |

| | | | |
|---|---|---|---|
| *Nannochloropsis gaditana* | Eustigmatophyte | EWM29759.1 | RSH1 |
| *Nannochloropsis gaditana* | Eustigmatophyte | EWM27974.1 | RSH4 |
| *Nannochloropsis gaditana* | Eustigmatophyte | EWM28606.1 | RSH4 |
| *Aureococcus anophagefferens* | Pelagophyte | XP_009034630.1 | RSH1 |
| *Aureococcus anophagefferens* | Pelagophyte | XP_009039283.1 | RSH1 |
| *Aureococcus anophagefferens* | Pelagophyte | XP_009039283.1 | RSH1 |
| *Aureococcus anophagefferens* | Pelagophyte | XP_009036518.1 | RSH4 |
| | **Chromalveolate/ Cryptophyta** | | |
| *Guillardia theta* | Cryptomonad | XP_005834725.1 | RSH4 |
| *Guillardia theta* | Cryptomonad | XP_005830237.1 | bacterial |
| | **Chromalveolate/ Haptophyte** | | |
| *Chrysochromulina* sp. | Prymnesiophyte | KOO29868.1 | RSH1 |
| *Chrysochromulina* sp. | Prymnesiophyte | KOO31885.1 | RSH1 |
| *Chrysochromulina* sp. | Prymnesiophyte | KOO35032.1 | RSH1 |
| *Chrysochromulina* sp. | Prymnesiophyte | KOO35751.1 | RSH1 |
| *Chrysochromulina* sp. | Prymnesiophyte | KOO21806.1 | RSH4 |
| *Chrysochromulina* sp. | Prymnesiophyte | KOO27310.1 | RSH4 |
| *Emiliania huxleyi* | Coccolithophore | XP_005769172.1 | - |
| *Emiliania huxleyi* | Coccolithophore | XP_005769365.1 | RSH1 |
| *Emiliania huxleyi* | Coccolithophore | XP_005777229.1 | RSH1 |
| *Emiliania huxleyi* | Coccolithophore | XP_005785512.1 | RSH1 |
| *Emiliania huxleyi* | Coccolithophore | XP_005759301.1 | RSH4 |
| *Emiliania huxleyi* | Coccolithophore | XP_005763353.1 | RSH4 |
| *Emiliania huxleyi* | Coccolithophore | XP_005779118.1 | RSH4 |
| *Emiliania huxleyi* | Coccolithophore | XP_005783324.1 | RSH4 |
| | **Chromalveolate/ Alveolate** | | |
| *Symbiodinium microadriaticum* | Dinoflagellate | OLP90598.1 | RSH1 |
| *Symbiodinium microadriaticum* | Dinoflagellate | OLP91394.1 | RSH1 |
| *Symbiodinium microadriaticum* | Dinoflagellate | OLP90600.1 | RSH4 |
| *Symbiodinium microadriaticum* | Dinoflagellate | OLP95483.1 | RSH4 |
| *Vitrella brassicaformis* | Chromerida | CEL95797.1 | - |
| | **Archaeplastida** | | |
| *Chlamydomonas reinhardtii* | Green alga | XP_001697429.1 | RSH2/3 |
| *Chlamydomonas reinhardtii* | Green alga | XP_001689858.1 | RSH2/3 |
| *Chlamydomonas reinhardtii* | Green alga | PNW86049.1 | RSH4 |
| *Chlorella variabilis* | Green alga | XP_005843741.1 | RSH1 |
| *Chlorella variabilis* | Green alga | XP_005849048.1 | RSH2/3 |
| *Chlorella variabilis* | Green alga | XP_005851775.1 | RSH4 |
| *Coccomyxa subellipsoidea* | Green alga | XP_005643555.1 | RSH1 |
| *Coccomyxa subellipsoidea* | Green alga | XP_005643557.1 | RSH2/3 |
| *Klebsormidium nitens* | Green alga | GAQ87841.1 | RSH1 |
| *Klebsormidium nitens* | Green alga | GAQ91082.1 | RSH4 |
| *Micromonas commode* | Green alga | XP_002505076.1 | RSH2/3 |

| | | | |
|---|---|---|---|
| *Micromonas pusilla* | Green alga | XP_003059515.1 | RSH1 |
| *Micromonas pusilla* | Green alga | XP_003060243.1 | RSH2/3 |
| *Ostreococcus lucimarinus* | Green alga | XP_001416591.1 | RSH4 |
| *Ostreococcus lucimarinus* | Green alga | XP_001420831.1 | RSH2/3 |
| *Ostreococcus lucimarinus* | Green alga | XP_001416051.1 | RSH2/3 |
| *Ostreococcus_tauri_* | Green alga | XP_003079611.1 | RSH1 |
| *Ostreococcus_tauri_* | Green alga | CEG01219.1 | RSH2/3 |
| *Ostreococcus_tauri_* | Green alga | OUS43685.1 | RSH4 |
| *Volvox carteri* | Green alga | XP_002958643.1 | RSH2/3 |
| *Volvox carteri* | Green alga | XP_002950935.1 | RSH4 |
| *Chondrus crispus* | Red alga | XP_005715321.1 | RSH1 |
| *Cyanidioschyzon merolae* | Red alga | XP_005538788.1 | RSH1 |
| *Cyanidioschyzon merolae* | Red alga | XP_005539556.1 | RSH1 |
| *Cyanidioschyzon merolae* | Red alga | XP_005536559.1 | RSH4 |
| *Cyanidioschyzon merolae* | Red alga | XP_005537243.1 | RSH4 |
| *Galdieria sulphuraria* | Red alga | XP_005705328.1 | RSH2/3 |
| *Galdieria sulphuraria* | Red alga | XP_005707095.1 | RSH2/3 |
| *Galdieria sulphuraria* | Red alga | XP_005705135.1 | RSH4 |
| *Galdieria sulphuraria* | Red alga | XP_005706615.1 | RSH4 |
| *Porphyra umbilicalis* | Red alga | OSX69462.1 | RSH1 |
| *Arabidopsis_thaliana* | Plant | ANM68011 | RSH1 |
| *Arabidopsis_thaliana* | Plant | AAF37282.1 | RSH2 |
| *Arabidopsis_thaliana* | Plant | AEE33053 | RSH3 |
| *Arabidopsis_thaliana* | Plant | OAP03147.1 | RSH4 |
| *Capsicum annuum* | Plant | NP_001311747.1 | RSH2 |
| *Cucumis melo* | Plant | XP_008439005.1 | RSH4 |
| *Hordeum vulgare* | Plant | BAJ88444.1 | RSH1 |
| *Hordeum vulgare* | Plant | BAJ95403.1 | RSH2/3 |
| *Hordeum vulgare* | Plant | BAJ99774.1 | RSH4 |
| *Ipomoea_nil* | Plant | XP_019163825.1 | RSH1 |
| *Ipomoea_nil* | Plant | ABV69554.1 | RSH2/3 |
| *Ipomoea_nil* | Plant | XP_019195966.1 | RSH4 |
| *Medicago truncatula* | Plant | KEH24410.1 | RSH1 |
| *Nicotiana tabacum* | Plant | XP_016480974.1 | RSH1 |
| *Nicotiana tabacum* | Plant | XP_016435380.1 | RSH2/3 |
| *Nicotiana tabacum* | Plant | XP_016454983.1 | RSH4 |
| *Oryza sativa* | Plant | XP_015632055.1 | RSH1 |
| *Oryza sativa* | Plant | XP_015611622.1 | RSH2/3 |
| *Oryza sativa* | Plant | XP_015644642.1 | RSH2/3 |
| *Oryza sativa* | Plant | XP_015637549.1 | RSH4 |
| *Oryza sativa* | Plant | BAF76773.1 | RSH4 |
| *Physcomitrella patens* | Plant | XP_024377135.1 | RSH1 |
| *Physcomitrella patens* | Plant | XP_024398657.1 | RSH1 |
| *Physcomitrella patens* | Plant | XP_024375791.1 | RSH2/3 |
| *Physcomitrella patens* | Plant | XP_024398683.1 | RSH2/3 |
| *Physcomitrella patens* | Plant | XP_024378912.1 | RSH2/3 |
| *Physcomitrella patens* | Plant | XP_024399053.1 | RSH2/3 |
| *Physcomitrella patens* | Plant | XP_024378514.1 | RSH2/3 |
| *Physcomitrella patens* | Plant | XP_024363061.1 | RSH4 |
| *Physcomitrella patens* | Plant | XP_024374499.1 | RSH4 |
| *Picea sitchensis* | Plant | ABR16061.1 | RSH1 |

| | | | |
|---|---|---|---|
| *Picea sitchensis* | Plant | ABK25186.1 | RSH4 |
| *Pisum sativum* | Plant | BAC97801.1 | RSH2/3 |
| *Populus trichocarpa* | Plant | XP_002314331.2 | RSH4 |
| *Populus trichocarpa* | Plant | XP_024441208.1 | RSH1 |
| *Populus trichocarpa* | Plant | XP_002298089.3 | RSH2/3 |
| *Ricinus communis* | Plant | XP_002523120.1 | RSH1 |
| *Ricinus communis* | Plant | XP_015576096.1 | RSH2/3 |
| *Ricinus communis* | Plant | XP_002519327.2 | RSH4 |
| *Selaginella moellendorffii* | Plant | XP_002976143.2 | RSH1 |
| *Selaginella moellendorffii* | Plant | XP_002973820.2 | RSH2/3 |
| *Selaginella moellendorffii* | Plant | XP_002977041.2 | RSH4 |
| *Sorghum bicolor* | Plant | XP_021302118.1 | RSH1 |
| *Sorghum bicolor* | Plant | XP_002460281.1 | RSH2/3 |
| *Sorghum bicolor* | Plant | XP_002463172.1 | RSH2/3 |
| *Sorghum bicolor* | Plant | XP_002439339.1 | RSH4 |
| *Vitis vinifera* | Plant | XP_010652206.1 | RSH1 |
| *Vitis vinifera* | Plant | CBI35865.3 | RSH2/3 |
| *Vitis vinifera* | Plant | XP_002268377.1 | RSH4 |
| *Zea mays* | Plant | XP_020399529.1 | RSH1 |
| *Zea mays* | Plant | ONM55312.1 | RSH2/3 |
| *Zea mays* | Plant | NP_001140933.1 | RSH4 |
| | **Rhizaria** | | |
| Paulinella chromatophora | Amoeba | YP_002048999.1 | |
| Paulinella micropora_ | Amoeba | APP88130.1 | |
| | **Prokaryote** | | |
| *Bacillus_subtilis_* | Bacterium | WP_087992279.1 | |
| *Brucella sp.* | Bacterium | WP_004683389.1 | |
| *Caldanaerobacter subterraneus* | Bacterium | KKC29792.1 | |
| *Caulobacter_vibrioides* | Bacterium | WP_010919427.1 | |
| *Dictyoglomus thermophilum* | Bacterium | WP_012547884.1 | |
| *Enterococcus faecalis_* | Bacterium | WP_104859299.1 | |
| *Escherichia coli* | Bacterium | AJF45176.1 | |
| *Marinithermus hydrothermalis* | Bacterium | WP_013704776.1 | |
| *Meiothermus silvanus* | Bacterium | WP_013156925.1 | |
| *Mesorhizobium sp.* | Bacterium | WP_010915098.1 | |
| *Neisseria meningitides* | Bacterium | WP_101294840.1 | |
| *Oceanithermus profundus* | Bacterium | WP_013457017.1 | |
| *Pseudomonas aeruginosa* | Bacterium | WP_096249453.1 | |
| *Rhizobiales* | Bacterium | WP_003527294.1 | |
| *Rhodobacter capsulatus* | Bacterium | WP_013069013.1 | |
| *Streptococcus pneumoniae* | Bacterium | WP_061632868.1 | |
| *Thermoanaerobacter sp.* | Bacterium | WP_013150236.1 | |
| *Thermomicrobium roseum* | Bacterium | WP_081433447.1 | |
| *Xylella fastidiosa* | Bacterium | WP_004089929.1 | |
| *Gloeobacter violaceus* | Cyanobacterium | WP_011142742.1 | |
| *Nostocaceae* | Cyanobacterium | WP_010995718.1 | |
| *Synechococcus* sp. WH 8102 | Cyanobacterium | WP_011129177.1 | |
| *Synechocystis* sp. PCC_6803 | Cyanobacterium | ALJ69438.1 | |
| *Thermosynechococcus vulcanus* | Cyanobacterium | BAY51563.1 | |

**Table S4  Additional domains found in RSH enzymes numbered in Fig.4**

| Code | Domain name | Abbreviation | Accession |
|---|---|---|---|
| 1 | EF-hand, calcium binding motif | EF-hand | Cd00051 |
| 2 | Tetratricopeptide repeat | TPR | COG0457 |
| 3 | Forked-associated domain | FHA | Pfam00498 |
| 4 | RNA recognition motif | RRM | Smart00360 |
| 5 | Ubiquitin domain of MoaD-like proteins | MoaD | Cd00754 |
| 6 | Major intrinsic protein | MIP | Cl00200 |
| 7 | Phosphopantetheinyl transferase | Sfp | COG2091 |
| 8 | Chlorophyll A-B binding protein | Chloroa_b-bind | Pfam00504 |
| 9 | RecX family | RECX | Cl00936 |
| 10 | Minor capsid protein VI | MCPVI | Pfam02993 |
| 11 | Endomucin Superfamily | EndoM | Cl25495 |
